# Neural filtering of physiological tremor oscillations to spinal motor neurons mediates short-term acquisition of a skill learning task

**DOI:** 10.1101/2023.07.20.549840

**Authors:** Hélio V. Cabral, Alessandro Cudicio, Alberto Bonardi, Alessandro Del Vecchio, Luca Falciati, Claudio Orizio, Eduardo Martinez-Valdes, Francesco Negro

## Abstract

The acquisition of a motor skill involves adaptations of spinal and supraspinal pathways to alpha motoneurons. In this study, we estimated the shared synaptic contributions of these pathways to understand the neural mechanisms underlying the short-term acquisition of a new force-matching task. High-density surface electromyography (HDsEMG) was acquired from the first dorsal interosseous (FDI; 7 males and 6 females) and tibialis anterior (TA; 7 males and 4 females) during 15 trials of an isometric force-matching task. For two selected trials (*pre-* and *post-skill* acquisition), we decomposed the HDsEMG into motor unit spike trains, tracked motor units between trials, and calculated the mean discharge rate and the coefficient of variation of inter-spike interval (CoV_ISI_). We also quantified the *post/pre* ratio of motor units’ coherence within delta, alpha, and beta bands. Improvements in force-matching were accompanied by a significant increase in the mean discharge rate and a decrease in CoV_ISI_ for both muscles. Moreover, the area under the curve within alpha band decreased by ∼22% and ∼13% for the TA and FDI muscles, respectively, with no changes in the delta or beta bands. These reductions correlated significantly with increased coupling between force/neural drive and target oscillations. These results suggest that the short-term acquisition of a new force-matching skill is mediated by the attenuation of tremor oscillations in the shared synaptic inputs. In other words, the central nervous system acts as a matched filter to modulate the synaptic weights of shared inputs and suppress neural components unrelated to the specific task. Supported by simulations, a plausible mechanism behind these alpha band reductions may involve spinal interneurons’ phase-cancelling descending oscillations.

## Introduction

Over the past three decades, the measurement of incremental motor skill acquisition has emerged as a relevant experimental paradigm for investigating the cognitive and neural processes underlying the learning of new motor abilities (Karni et al., 1998; Ungerleider et al., 2002). Initially, the new movement is produced with varying accuracy, but with repetition and practice, the central nervous system refines the movement, resulting in effortless and precise execution (Willingham, 1998). Consequently, the learning of a new motor task entails structural and functional alterations in the supraspinal and spinal pathways (i.e., neural plasticity) to accommodate the acquisition and retention of skilled motor behaviors (Ungerleider et al., 2002; Dayan and Cohen, 2011). Indeed, compelling evidence from studies on primates (Plautz et al., 1995; Nudo et al., 1996) and non-primates (Kleim et al., 1998) animals have demonstrated that, following motor skill training, there is an expansion of the cortical representations related to the trained task. Similar neural adaptations have been observed in humans in studies using non-invasive imaging and neurophysiological techniques (Karni et al., 1995; Pascual-Leone et al., 1995; Perez et al., 2004; Jensen et al., 2005). Specifically, the acquisition of new motor skills has been shown to significantly increase cortical representation (Karni et al., 1995; Pascual-Leone et al., 1995; Pascual-Leone et al., 1999) and corticospinal excitability (Perez et al., 2004; Jensen et al., 2005) of the muscles involved in the training task. While there is greater consensus regarding the effects of motor skill learning on supraspinal adaptations in humans, the dynamics of plastic changes in synaptic connectivity to spinal motor neurons remain relatively unexplored in the literature.

Many daily activities require the rapid acquisition of motor tasks that demand precise control of fine movements within short time intervals (Johansson and Westling, 1988). As the generation of precise movements relies on the accurate modulation of muscle forces (Johansson and Westling, 1988; Kumar et al., 2017), specific control strategies are necessary during motor skill learning to overcome the inherent variability of the neural pathways innervating muscles. Interestingly, previous research has demonstrated that the motor neuron pool behaves as a very selective filter, eliminating components of synaptic input that are not common to all motor neurons (Negro and Farina, 2011a; Farina et al., 2014; Negro et al., 2016a). Additionally, it has also been observed that the muscle itself acts as a spatial and temporal smoothing filter of the neural drive (Bawa and Stein, 1976; Baldissera et al., 1998), further minimizing the effects of independent sources of neural synaptic noise across the alpha motor neuron pools. Collectively, these investigations indicate that only the low-frequency components of the synaptic inputs widely shared across the motor neuron pools are represented in the force output (Enoka and Farina, 2021). In voluntary tasks, these shared synaptic inputs consist of task-related oscillations, which determine the precise command for optimal force generation (e.g., control input signals from cortico-spinal pathways), as well as task-unrelated oscillations. These task-unrelated oscillations comprise both voluntary components (reflecting errors in task performance) and involuntary components (e.g., physiological tremor) that act as noise, thereby reducing the precision of the task. Thus, the process of learning new motor tasks involving precise force generation should require minimizing these task-unrelated oscillations, whether voluntary or involuntary, to maximize the representation of shared synaptic input related to optimal control in the force output (i.e., task-related oscillations).

Despite extensive research exploring the effect of motor learning on reducing errors in task performance (Ungerleider et al., 2002; Dayan and Cohen, 2011), no study has yet experimentally investigated whether the acquisition of a new motor skill involves a reduction in the involuntary components of shared synaptic input. Particularly intriguing is the observation that during steady contractions, the force output exhibits an involuntary oscillation at ∼10 Hz (alpha band), commonly associated with physiological tremor (McAuley and Marsden, 2000). The origin of these rhythmic fluctuations has been attributed to cortical pathways (Marsden et al., 1967; McAuley et al., 1997; Evans and Baker, 2003) and/or peripheral pathways, likely originating from the Ia afferent feedback loop (Halliday and Redfearn, 1956; Lippold, 1971). If indeed the acquisition of new motor skills is mediated by reductions in physiological tremor, specific cortical and/or peripheral neural mechanisms must emerge during the skill learning task to reduce alpha band oscillations in the shared synaptic input to motor neurons. For instance, previous data in macaque monkeys have shown that spinal interneurons could phase-cancel ∼10Hz cortical inputs to motor neurons, which would be beneficial to decrease tremor and improve movement precision (Williams et al., 2010; Koželj and Baker, 2014). Therefore, it is possible that during motor skill learning, the gain of this spinal interneurons filter could be upregulated, thereby increasing the cancelation. On the other hand, compelling evidence has demonstrated that changes in the gain of afferent feedback loop may directly modulate tremor inputs (Cresswell and Löscher, 2000; Christakos et al., 2006; Laine et al., 2016), presenting another potential mechanism that could be involved in the acquisition of a skill learning task. Even though both mechanisms appear intuitively reasonable, no study to date has explored whether either of these mechanisms plays a role during the acquisition of a skill-learning task.

The present study aimed to investigate whether the short-term acquisition of a new force-matching skill is mediated by alterations in the shared synaptic input to spinal motor neurons, particularly in the physiological tremor band (alpha band; 5-15 Hz). We hypothesized that the acquisition of a new motor skill in an individual muscle would require specific adaptations in the neural pathways to the motor neuron pool of that muscle, ultimately reducing the contributions of shared synaptic oscillations unrelated to the task (i.e., alpha band oscillations).

We experimentally tested this hypothesis by decomposing high-density surface electromyograms from the first dorsal interosseous (FDI) and tibialis anterior (TA) muscles during 15 trials of a complex, isometric force-matching task (force-matching skill acquisition). In addition, we simulated a population of motor neurons receiving both common and independent inputs to elucidate the potential neural mechanisms underlying the experimental results. Specifically, we simulated two scenarios: filtering of alpha oscillations by spinal interneuron circuits (scenario A) and filtering of alpha oscillations in the Ia afferent feedback loop (scenario B). We then compared the results of the two scenarios with the experimental results, aiming to determine which scenario best explains the observed experimental outcomes.

## Results

### Changes in force-matching and force power spectrum with skill acquisition

To assess improvements in force-matching during the skill acquisition task, we compared the root-mean-square error (RMSE) and cross-correlation peak values between the force and target signals in the first two trials and the last two trials out of the 15. **Figure 1A** shows the fluctuations in dorsiflexion isometric force produced by a representative participant for these trials, where the yellow traces represent the first two trials, and the green traces refer to the last two. Improvements in force-matching are evident across trials, which is visually confirmed by the greater overlap between the force and the target template when comparing the final two with the initial two trials. Indeed, for both TA and FDI muscles, there were significant differences in RMSE (*P* < 0.001 for both muscles, Friedman test) and cross-correlation peak values (*P* < 0.001 for both muscles, Friedman test) among trials. Specifically, for both muscles, RMSE values were significantly lower in the last two trials compared with the first two (*P* < 0.006 for all, Bonferroni post-hoc test; **Figures 1B** and **1C**). Moreover, for both muscles, cross-correlation peak values obtained in the last two trials were significantly greater than in the first two (*P* < 0.006 for all, Bonferroni post-hoc tests). No significant differences were found in RMSE and cross-correlation within both the initial two trials and the final two trials (*P* > 0.128 for all, Bonferroni post-hoc tests). Given this lack of difference and with the aim of maximizing the number of tracked motor units between trials (see *Methods* section), for all subsequent analyses, we selected one trial from the first two (the one with the highest RMSE) and one trial from the last two (the one with the lowest RMSE) to represent the *pre-* and *post-skill acquisition* trials.

**Figure 1:**
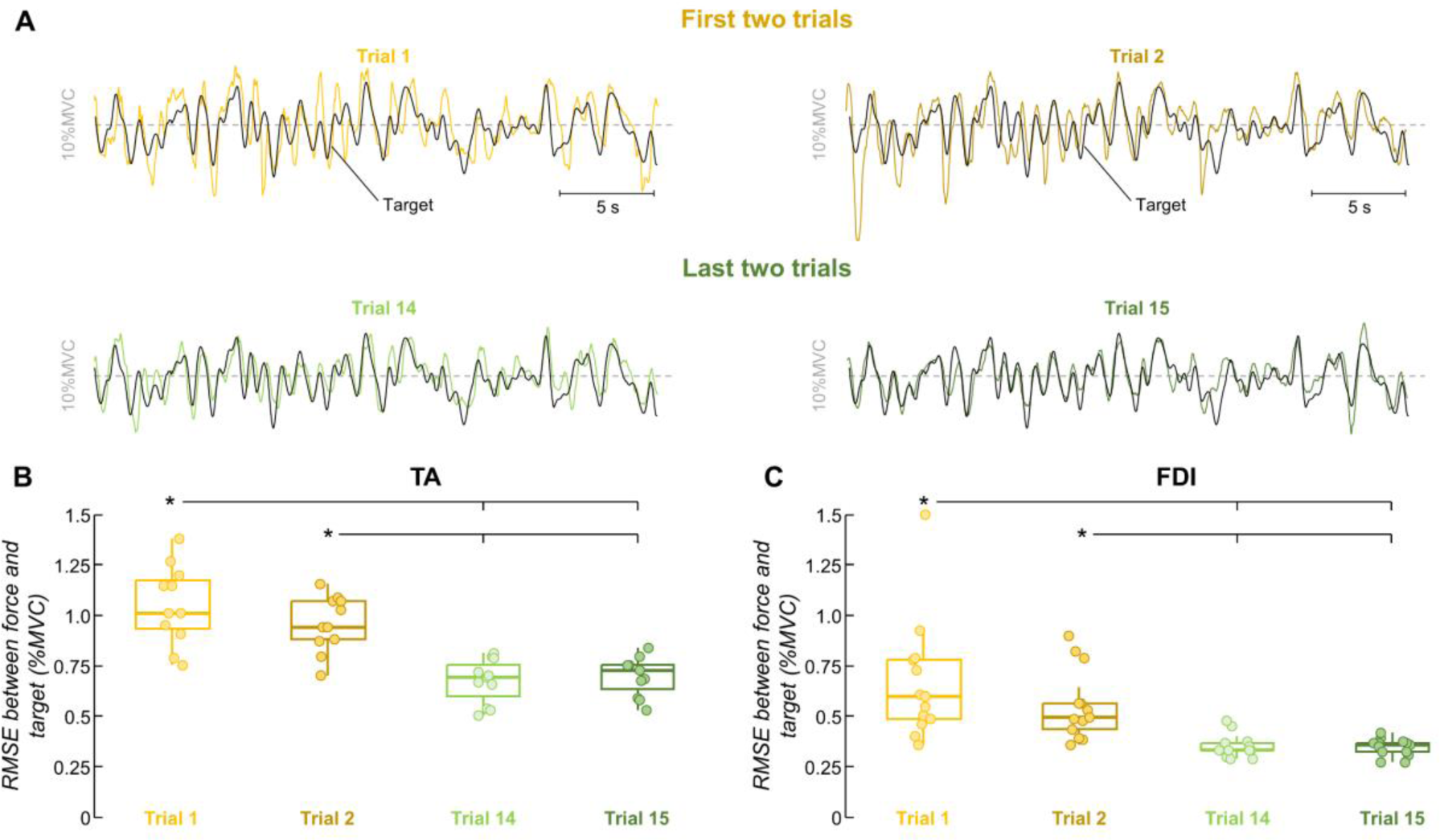
Performance results. Four (first two and last two) out of the 15 trials were used for each participant to assess improvements in force-matching. A, Representative comparison between the force and target during the skill acquisition task. The yellow lines indicate dorsiflexion isometric forces produced by a participant for the first two trials, and the green lines for the last two trials. The black line indicates the target. B-C, Group results of root-mean-square-error (RMSE) between the force and target for the tibialis anterior (TA) muscle (B) and the first dorsal interosseous (FDI) muscle (C). Circles identify individual participants. Horizontal traces, boxes, and whiskers denote the median value, interquartile interval, and distribution range. *P<0.05.

In **Figure 2A**, it is possible to see the two trials that were chosen to represent the *pre-* and *post-skill acquisition* trials for the same participant in **Figure 1A**. There is a decrease of ∼43% in RMSE (from 1.16 to 0.66 %MVC) and an increase of ∼88% in cross-correlation peak value (from 0.41 to 0.78) between *pre-* and *post-acquisition* trials. To assess whether the force steadiness changed with the force-matching skill acquisition, we quantified the coefficient of variation of force for the two selected trials. For both TA and FDI muscles, there were significant differences in the coefficient of variation of force between trials (**Figure 2B**). Specifically, the coefficient of variation of force values obtained for the *post-skill acquisition* trial were significantly lower than *pre-skill acquisition* trial for both TA (*pre*: 11.77 ± 1.57%, *post*: 9.51 ± 0.83%; *P* = 0.005; Wilcoxon signed-rank test) and FDI (*pre*: 14.13 ± 6.18%, *post*: 8.65 ± 0.93%; *P* < 0.001, Wilcoxon signed-rank test). We also examined how mean force power changed during the learning task, separately for the delta (1-5 Hz), and alpha (5-15 Hz) bands (the frequency bandwidth of the force signal). **Figure 2C** illustrates the power spectrum of force signals depicted in **Figure 2A**. It is visually evident for this representative participant that there was a reduction in the power spectrum of force between *pre-* and *post-skill acquisition* trials for both frequency bandwidths. This reduction in the mean force power with the force-matching skill acquisition was statistically confirmed for both TA (*P* < 0.003 for both delta and alpha bands, Wilcoxon signed-rank test; **Figure 2D**) and FDI (*P* < 0.002 for both delta and alpha bands, Wilcoxon signed-rank test; **Figure 2E**) muscles.

**Figure 2:**
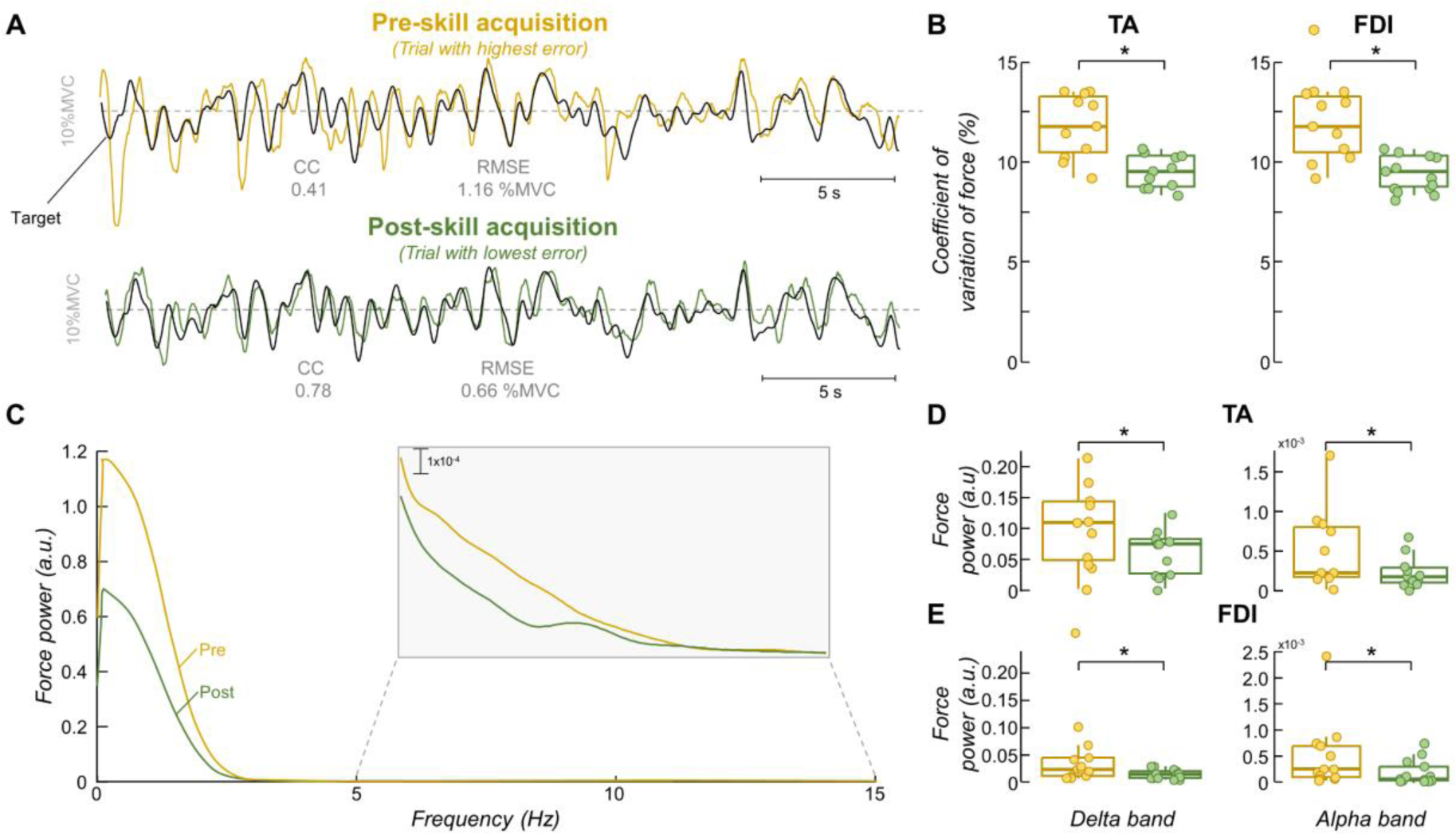
Force steadiness and force power spectrum results. Two trials were selected for each participant to represent the pre- and post-skill acquisition trials. A, Representative comparison between the force and target for these two trials, where the yellow and green lines indicate the pre- and post-skill acquisition trials, respectively. The black line indicates the target. B, Group results of coefficient of variation of force (force steadiness). C, Power spectrum of force signals depicted in A. The grey box shows a zoom in the alpha band (5-15 Hz). D-E, Group results of mean force power the tibialis anterior (TA) muscle (D) and the first dorsal interosseous (FDI) muscle (E). Circles identify individual participants. Horizontal traces, boxes, and whiskers denote median value, interquartile interval, and distribution range. *P<0.05.

### Changes in motor unit discharge properties with skill acquisition

In order to evaluate changes in motor unit discharge properties (i.e., mean discharge rate and coefficient of variation of inter-spike interval (COV_ISI_)) with the force-matching skill acquisition, we decomposed high-density surface electromyograms (HDsEMG) signals into motor unit spike trains. Note that we tracked motor units *pre-* and *post-skill acquisition* to ensure the same motor units were compared between trials (see *Methods* section for further details). For the TA muscle, we identified a total of 166 and 198 motor units for *pre-* and *post-skill acquisition* trials, respectively, and were able to track 138 motor units (13 ± 7 motor units per participant). For the FDI muscle, instead, we identified a total of 105 and 127 motor units for *pre-* and *post-skill acquisition* trials, respectively, from which 62 were tracked across trials (5 ± 2 motor units per participant).

For both muscles, a significant effect of trial (*pre-* vs *post-skill acquisition*) was found on the mean discharge rate and COV_ISI_ values of matched motor units. Specifically, for the TA muscle, the mean discharge rate significantly increased from 10.9 [9.95, 11.9] to 11.9 [10.92, 12.9] pps between *pre-* and *post-skill acquisition* trials (*F* = 23.083, *P* < 0.001, Linear mixed models (LMM); **Figure 3A**). Conversely, the COV_ISI_ significantly decreased from 48.0 [42.9, 53.2] to 31.4 [26.3, 36.3] % with the force-matching skill acquisition (*F* = 22.986, *P* < 0.001, LMM; **Figure 3C**). Similar results were observed for the FDI muscle, with the values of mean discharge rate significantly increasing from 11.6 [10.5, 12.7] to 12.3 [11.3, 13.4] pps (*F* = 5.166, *P* = 0.025, LMM; **Figure 3B**), and the values of COV_ISI_ significantly decreasing from 46.6 [39.4, 53.8] to 35.4 [28.1, 42.6] % (*F* = 8.952, *P* = 0.003, LMM; **Figure 3D**) between *pre-*and *post-skill acquisition* trials.

**Figure 3:**
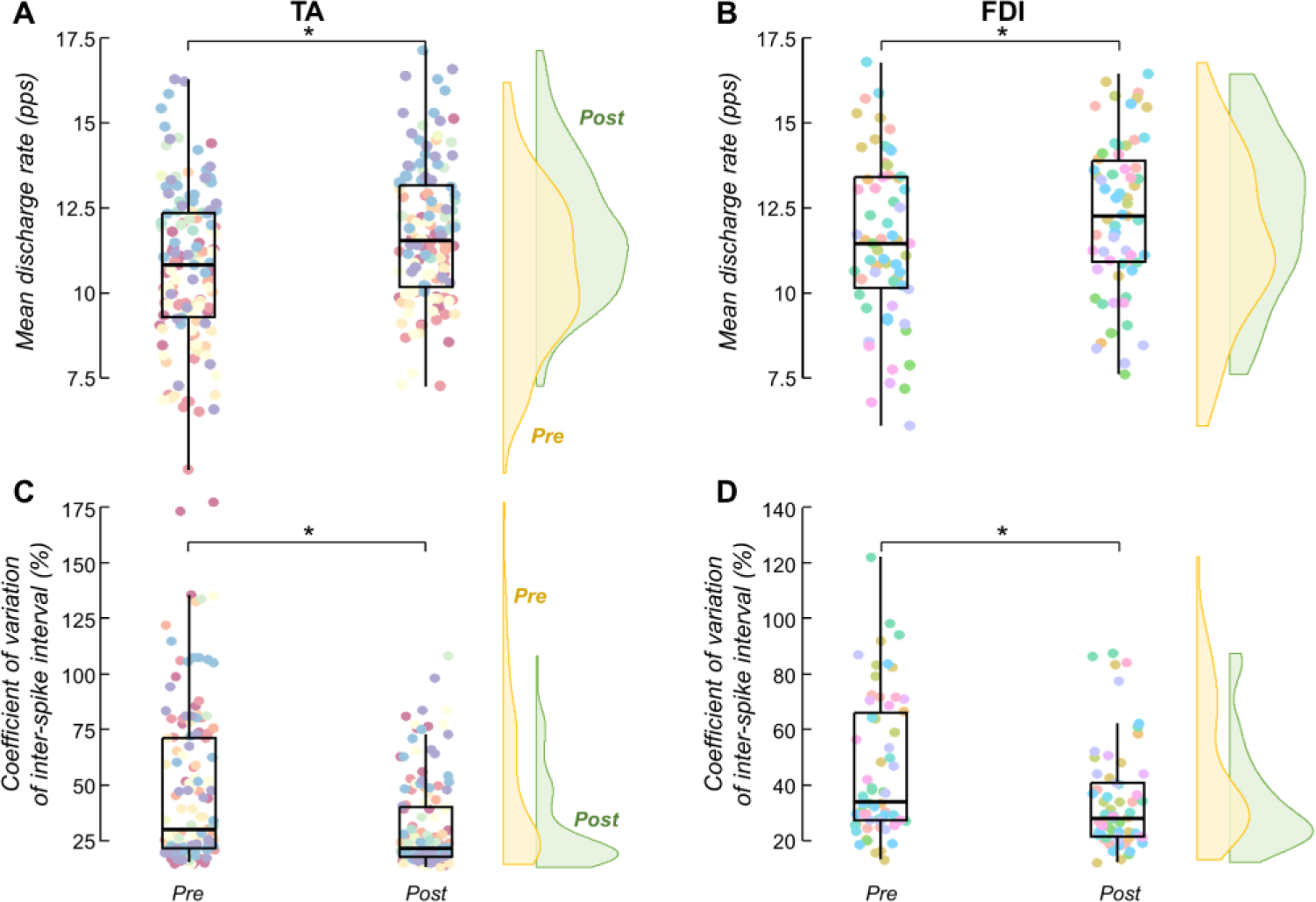
Mean discharge rate and discharge variability results. A-B, mean discharge rate results of matched motor units between pre- and post-skill acquisition trials for the tibialis anterior (TA) muscle (A) and the first dorsal interosseous (FDI) muscle (B). C-D, coefficient of variation of inter-spike interval results of matched motor units between pre- and post-skill acquisition trials for the TA (C) and FDI (D) muscles. Each circle identifies a matched motor unit between trials. Each color of the circles corresponds to a specific participant. Horizontal traces, boxes, and whiskers denote median value, interquartile interval, and distribution range. Density curves of the data are represented on the right side of each panel by half-violin plots (yellow for pre-skill acquisition and green for post-skill acquisition). Note that density curves can be used to visually compare differences between pre- and post-skill acquisition trials. *P<0.05.

### Changes in common synaptic input with skill acquisition

In order to assess changes in common synaptic input between *pre-* and *post-skill acquisition* trials, we used coherence analysis between the spike trains of matched motor units. We specifically quantified the ratio (*post/pre*) of the area under the curve of z-coherence within delta (1-5 Hz), alpha (5-15 Hz), and beta (15-35 Hz) bands. The coherence analysis was performed in 10 and 9 participants for TA and FDI muscles, respectively, as only three or fewer motor units were matched for participant #7 in TA, and participants #2, #8, #11, and #13 in FDI. **Figures 4A** and **4B** display the pooled z-coherence for all participants. When comparing *pre-* and *post-skill acquisition* trials, for both TA and FDI muscles there is a clear reduction in the area under the curve in the alpha band (grey area in **Figure 4A** and **4B**). Indeed, for the TA muscle, there was a significant median reduction of ∼22% in the area under the curve within the alpha band between *pre-* and *post-skill acquisition* trials (*P* = 0.014, one-sample Wilcoxon signed-rank test; **Figure 4C**), which was not observed either for delta or beta bands (*P* > 0.322 for both, one-sample Wilcoxon signed-rank tests). Similarly, for the FDI muscle, the area under the curve within the alpha band significantly decreased by a median of ∼13% with the force-matching skill acquisition (*P* = 0.008, one-sample Wilcoxon signed-rank test; **Figure 4D**) but did not significantly change for delta or beta bands (*P* > 0.074 for both, one-sample Wilcoxon signed-rank tests).

**Figure 4:**
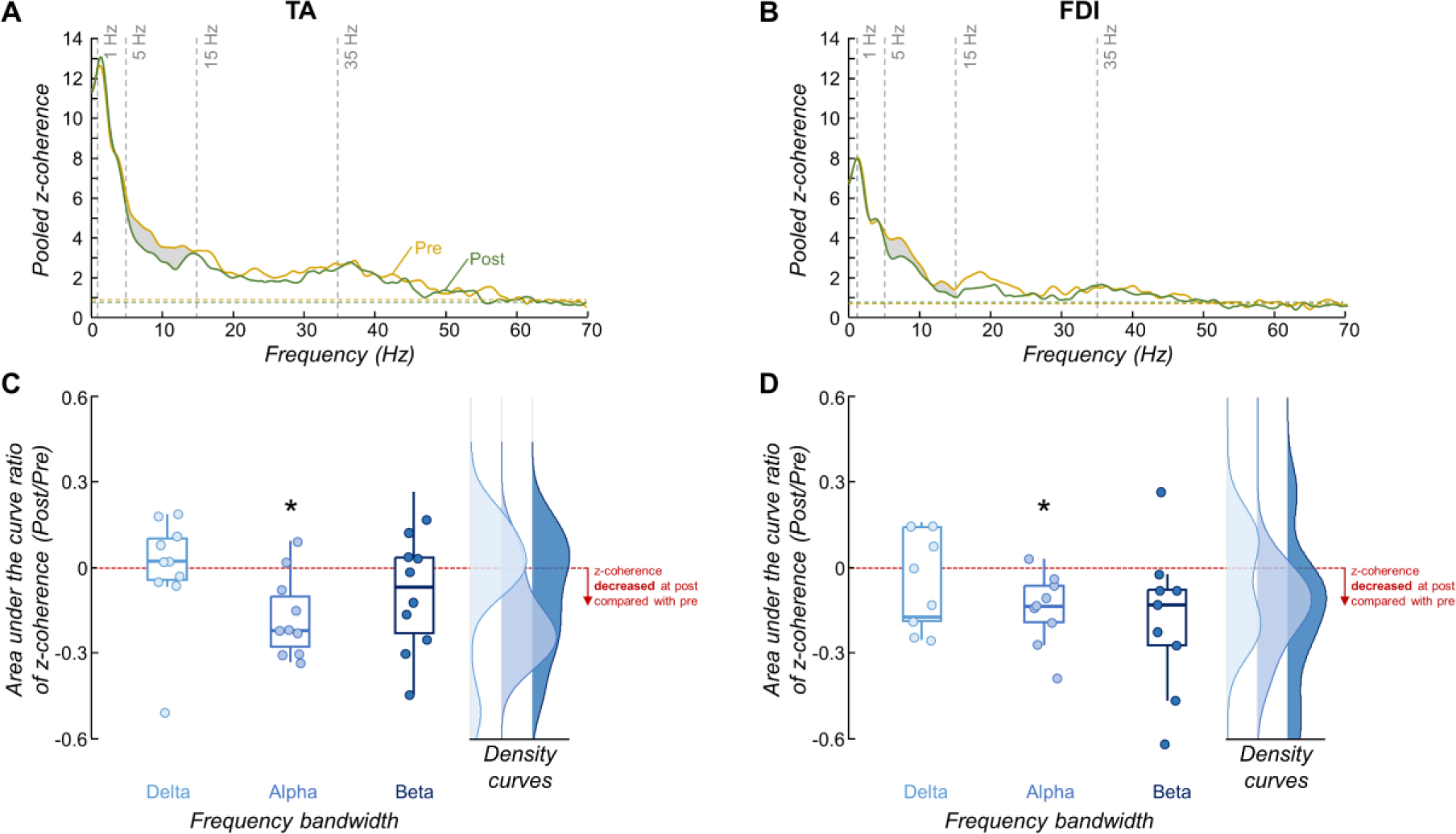
Motor unit coherence results. A-B, pooled z-coherence profiles considering all participants for the tibialis anterior (TA) muscle (A) and first dorsal interosseous (FDI) muscle (B) (yellow for pre-skill acquisition and green for post-skill acquisition). The horizontal dashed line indicates the confidence level. Vertical dashed lines highlight the three frequency bandwidths analyzed: delta (1-5 Hz), alpha (5-15 Hz), and beta (15-35 Hz) bands. Grey areas denote statistical differences in the area under the curve between pre- and post-skill acquisition trials. C-D, group results of the area under the curve ratio of coherence for the TA (C) and FDI (D) muscles. Circles identify individual participants. Horizontal traces, boxes, and whiskers denote median value, interquartile interval, and distribution range. Density curves of the data are represented on the right side of each panel by half-violin plots. *P<0.05.

To examine whether the absence of statistical differences within the beta band was due to the number of motor units utilized in the cumulative spike trains (CST), we conducted an additional analysis using the TA motor unit data (for detailed information, please refer to the *Methods* section). In brief, we calculated z-coherence between two CSTs, incrementally increasing the number of motor units in each CST from 1 to 13 (half of the maximal number of matched motor units across all participants). We then computed the area under the curve ratio (*post/pre*) of z-coherence within the beta band. If our results within the beta band were influenced by the number of motor units used in the CSTs, we would expect to observe a significant deviation from zero in this ratio curve as the number of motor units increased. Instead, we observed that the results of area under the curve ratio of z-coherence within the beta band were not influenced by the number of motor units used in the CSTs, which was confirmed statistically using the one-sample Statistical Parametric Mapping (*P* > 0.05).

### Changes in coherence between force/neural drive and target with skill acquisition

To evaluate changes in the linear coupling between oscillations in force/neural drive and oscillations in the target template between *pre-* and *post-skill acquisition* trials, we calculated the z-coherence within delta band (the frequency bandwidth of the target) between the target and the force, and between the target and the neural drive to the muscles (i.e., CST). **Figure 5** displays the pooled z-coherence between force and target, and between CST and target, for both TA (top) and FDI (bottom) muscles. In all cases, there was a clear increase in the area under the curve of z-coherence between *pre-* and *post-skill acquisition* trials. Indeed, significant increases in the area under the curve ratio of target-force z-coherence (*P* < 0.001 for both muscles, one-sample Wilcoxon rank-signed test; **Figure 7A**) and target-CST z-coherence (*P* < 0.004 for both muscles, one-sample Wilcoxon rank-signed test; **Figure 7B**) were observed between *pre-* and *post-skill acquisition* trials.

**Figure 5:**
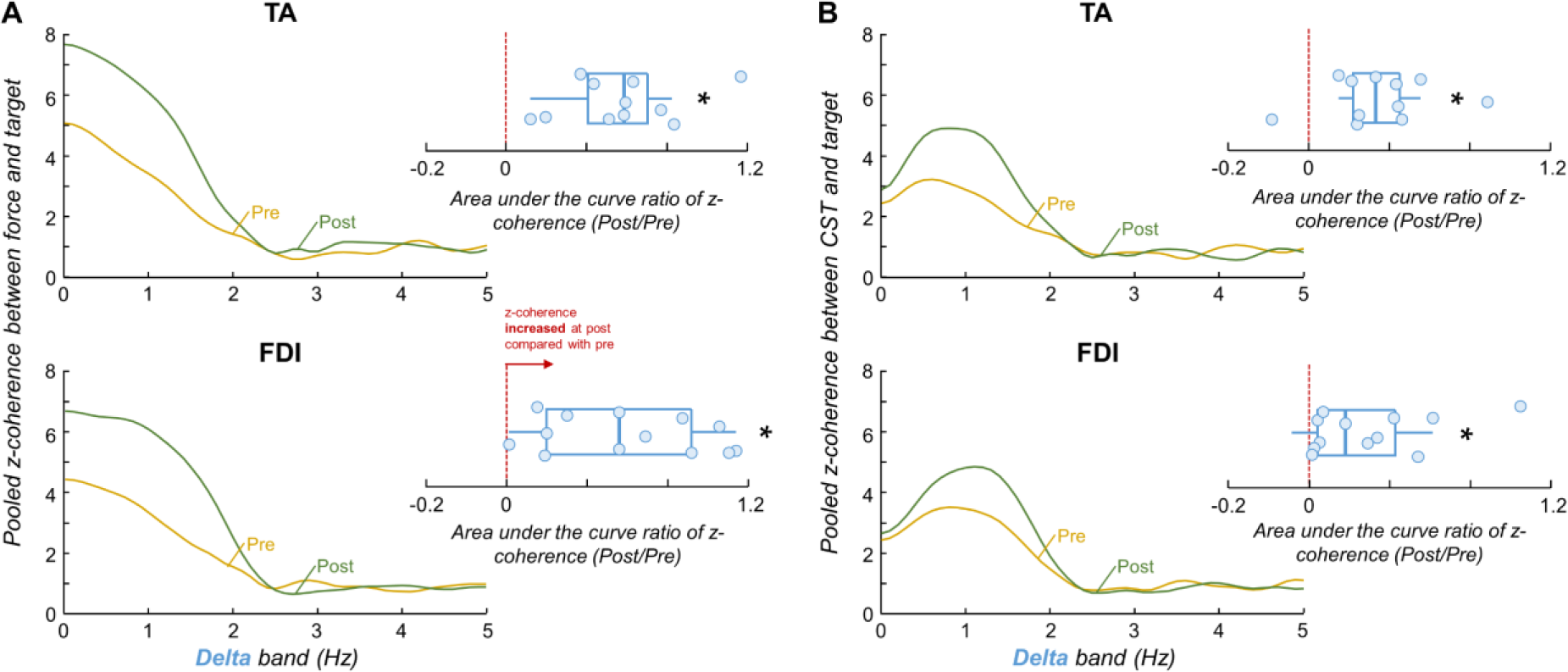
Coherence between force/neural drive and target results. Pooled z-coherence profiles between force and target (A) and cumulative spike train (CST) and target (B) for the tibialis anterior (TA) muscle (top) and first dorsal interosseous (FDI) muscle (bottom). Yellow and green lines indicate the pre-skill acquisition and post-skill acquisition trials, respectively. Note that the frequency bandwidth analyzed was only the delta band (the frequency bandwidth of the target). Blue boxplots show the group results of the area under the curve ratio of z-coherence. Circles identify individual participants. Horizontal traces, boxes, and whiskers denote median value, interquartile interval, and distribution range. *P<0.05.

**Figure 6:**
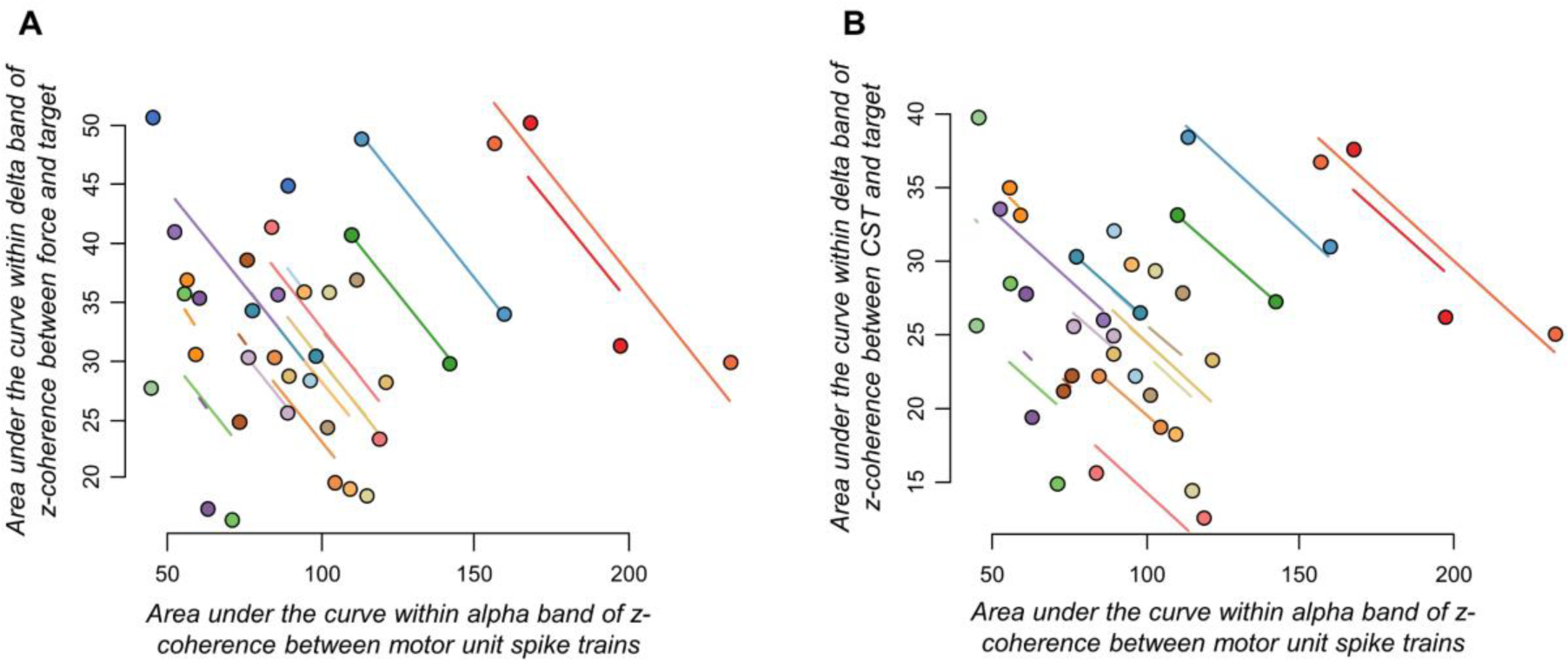
Correlation results. Repeated measures correlations between changes in motor unit coherence within the alpha band and coherence between force and target (A), as well as between motor unit coherence within the alpha band and coherence between cumulative spike train (CST) and target (B). For this analysis, data from tibialis anterior and first dorsal interosseous muscles were pooled together.

**Figure 7:**
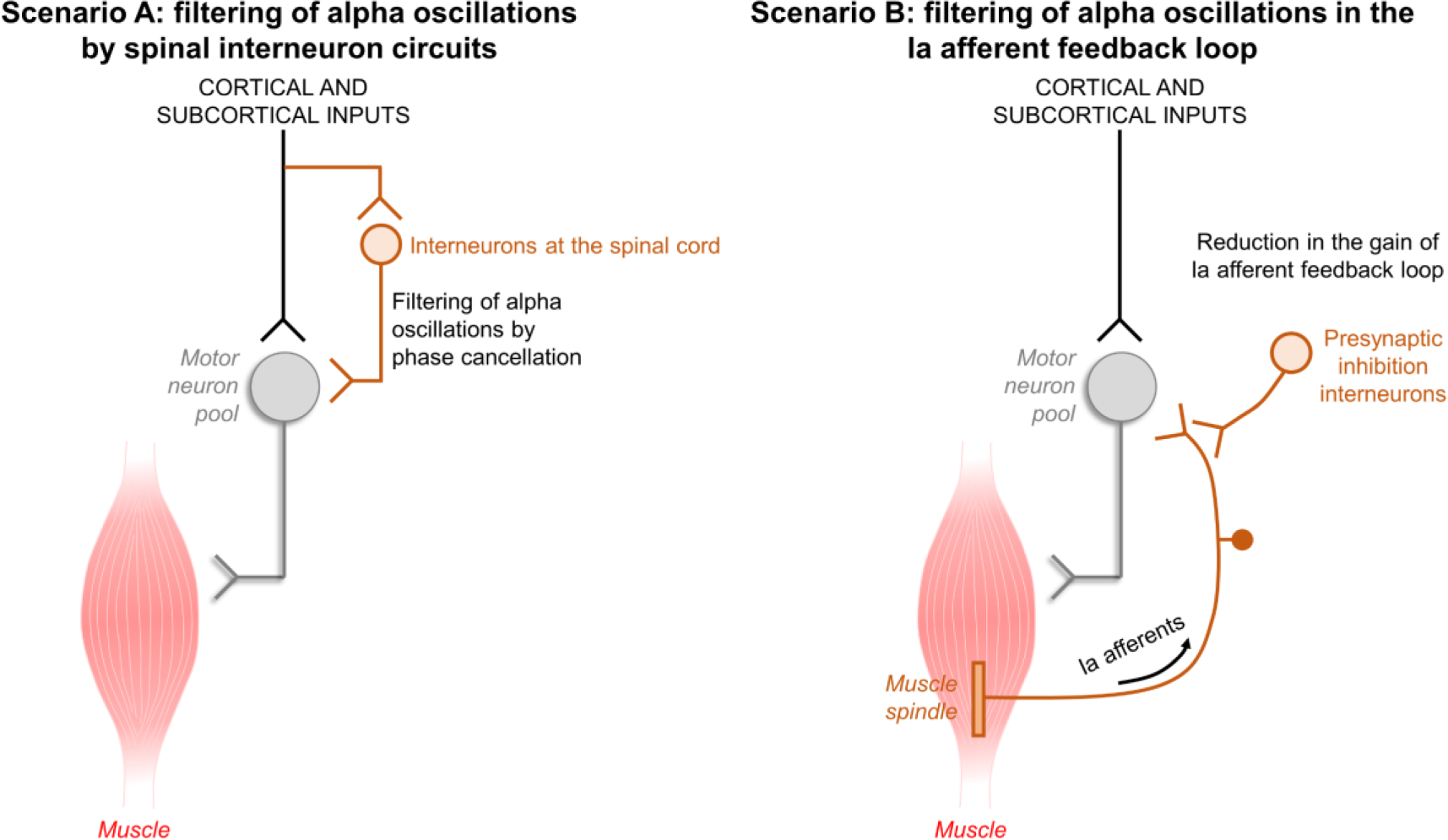
Simulated scenarios to investigate neural mechanisms underlying the experimental results. Two different scenarios were simulated to investigate potential mechanisms that could explain the observed changes between pre- and post-skill acquisition. In scenario A (left panel), we hypothesized that decreases in alpha band with the acquisition of the force-matching skill could be explained by spinal interneurons phase-cancelling central oscillatory inputs in the alpha frequency range. In scenario B (right panel), we hypothesized that reductions in alpha band with the force-matching skill acquisition could be explained by increases in presynaptic inhibition of Ia afferent feedback into the motor neuron pool. Details about how we simulated these scenarios are provided in the Methods section.

To investigate whether reductions in the motor unit z-coherence within the alpha band between trials (**Figure 4**) were correlated with changes in target-force and target-CST coherence between trials (**Figure 5**), we used repeated measures correlations. For this analysis, the data from TA and FDI muscles were pooled together. **Figure 6** shows a significant inverse association between alpha band coherence and target-force coherence (**Figure 6A**; *P* = 0.002), as well as between alpha band coherence and target-force coherence (**Figure 6B**; *P* = 0.004).

### Neural mechanisms underlying the short-term acquisition of a skill task

To explore the neural mechanisms underlying the observed changes between *pre-* and *post-skill acquisition*, we simulated a population of motor neurons receiving common and independent inputs and the sequence of events from the excitation of these motor neurons to the generation of isometric force output. Specifically, two different scenarios were simulated (refer to *Methods* section for further details). In scenario A, we hypothesized that decreases in alpha band with the acquisition of the force-matching skill could be explained by spinal interneurons phase-cancelling central oscillatory inputs in the alpha frequency range (**Figure 7A**). In scenario B, the reductions in alpha band with the force-matching skill acquisition could be explained by increases in presynaptic inhibition of Ia afferent feedback into the motor neuron pool (**Figure 7B**). We calculated the simulated force output power spectrum; the ratio (*post/pre*) of the area under the curve of motor unit z-coherence within delta (1-5 Hz), alpha (5-15 Hz) and beta (15-35 Hz) bands; and the area under the curve ratio (post/pre) of z-coherence between simulated force/CST and the target template. We then compared the results of the different scenarios with the experimental results to explore which simulated scenario aligns more closely with the observed experimental outcomes.

Results of scenario A were found to align more closely with the observed experimental outcomes, but only when we slightly increased the beta band input to the motor neuron pool in the *post-skill acquisition* model compared with *pre-skill acquisition* model. Consistent with the experimental results, there was a reduction in the power spectrum of force between *pre-* and *post-skill acquisition* models for both delta and alpha bands (P < 0.05 for both, Wilcoxon signed-rank test). **Figure 8A** displays the pooled z-coherence of simulated motor units for the ten realizations of the best fitting scenario (scenario A), showing a clear reduction within alpha band between *pre-* (yellow line) and *post-skill acquisition* (green line) models. Indeed, the area under the curve of coherence within the alpha band significantly decreased between *pre-* and *post-skill acquisition* models (*P* < 0.002, one-sample Wilcoxon signed-rank test; **Figure 8B**) but did not significantly change either for delta or beta bands (*P* > 0.274 for both; one-sample Wilcoxon signed-rank tests; **Figure 8B**). Moreover, there were an increase in the area under the curve of simulated target-force z-coherence (*P* = 0.002, one-sample Wilcoxon rank-signed test; **Figure 8C**) and simulated target-CST z-coherence (*P* = 0.002, one-sample Wilcoxon rank-signed test; **Figure 8B**) between *pre-* and *post-skill acquisition* models. In contrast, the simulation results of scenario B did not match the experimental results. In this scenario, there was a significant increase in the power spectrum of force within delta band (*P* = 0.002, Wilcoxon signed-rank test), as well as a significant increase in the area under the curve of coherence within the delta band (*P* = 0.002, one-sample Wilcoxon signed-rank test) between *pre-* and *post-skill acquisition* models, which were not observed in the experimental outcomes.

**Figure 8:**
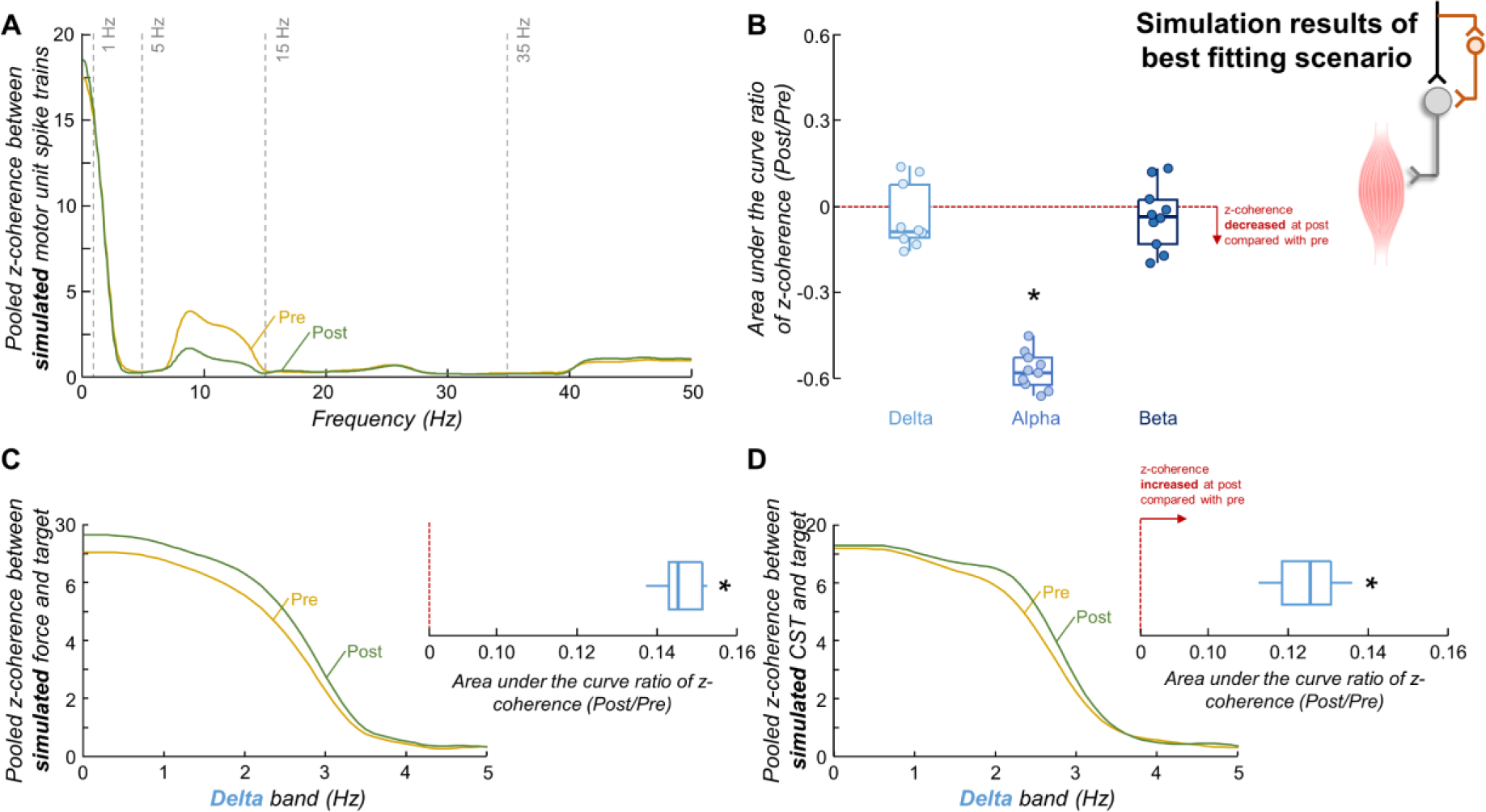
Simulation results of best fitting scenario. A, pooled z-coherence profiles considering all realizations. Vertical dashed lines highlight the three frequency bandwidths analyzed: delta (1-5 Hz), alpha (5-15 Hz), and beta (15-35 Hz) bands. B, group results of the area under the curve ratio of coherence. Circles identify individual simulation realizations. Horizontal traces, boxes, and whiskers denote median value, interquartile interval, and distribution range. C-D, pooled z-coherence profiles between simulated force and target (C) and between simulated cumulative spike train (CST) and target (D). Note that the frequency bandwidth analyzed was only the delta band (the frequency bandwidth of the target). Blue boxplots show the results of the area under the curve ratio of z-coherence for all simulation realizations. In panels A, C and D, yellow and green lines indicate the pre-skill acquisition and post-skill acquisition models, respectively. *P<0.05.

## Discussion

In this study, we investigated whether short-term learning of a complex, isometric force-matching task is mediated by specific adaptations in the shared synaptic inputs to alpha motor neuron pools of the TA and FDI muscles. Our experimental findings revealed that both muscles exhibited improvements in force-matching as skill acquisition progressed, accompanied by a reduction in coherent oscillations across motor neuron spike trains unrelated to the required force fluctuations (i.e., alpha or tremor band oscillations). Importantly, these reductions in alpha band with the force-matching skill acquisition correlated significantly with an increased coupling between force/neural drive and target oscillations. Based on simulations, our findings further indicate that the potential neural mechanism underlying decreases in alpha band with the acquisition of the force-matching skill is related to spinal interneurons phase-cancelling central oscillatory inputs in the alpha frequency range. As discussed below, these outcomes suggest that the acquisition of a new force-matching task involves specific changes in spinal neural circuitry that behave as a matched neural filter, ultimately minimizing shared noise components unrelated to the intended task.

Traditionally, motor sequence learning, which assesses the incremental acquisition of sequential motor skills, has been used as experimental paradigm to investigate neuroplasticity underlying skilled task acquisition (Karni et al., 1998; Ungerleider et al., 2002). Psychophysiological evidence has revealed that two main stages are involved in this paradigm: a fast stage characterized by considerable performance improvements within a single session, and a slow stage where further but quantitatively smaller improvements can be observed across multiple sessions (Ungerleider et al., 2002; Dayan and Cohen, 2011). Consistent with previous research (Knight and Kamen, 2004; Perez et al., 2005), our study focused on the fast-learning stage of incremental motor skill acquisition, as we aimed to investigate the rapid acquisition of a fine control task involving muscle force modulation. Thus, ensuring that the observed neural plastic changes in this study were specifically related to the force-matching skill task, and did not arise from overall motor activity or other confounding factors, is a necessary step for interpreting our main results. The median reductions in RMSE of ∼40% for both the TA and FDI observed after 15 repetitions of the task (**Figure 1**), along with cross-correlation increases of ∼35% for both muscles, indicate that the chosen task and number of trials were sufficiently challenging to enable skill acquisition. Previous experiments using a similar task involving index finger abduction demonstrated comparable RMSE improvements (∼50%) between force and target over the same number of repetitions (Knight and Kamen, 2004). Taking these factors into consideration, it is plausible to assume that the alterations identified in motor unit discharge characteristics and shared synaptic inputs are indeed associated with the intended force-matching task.

Due to the low-pass and amplification characteristics of the motor neuron pool, experimental and simulated data have extensively demonstrated that only the low-frequency components of the synaptic inputs largely shared across the motor neuron pool are represented in the generated muscle force (Negro et al., 2009; Negro and Farina, 2011b, a; Farina et al., 2014; Negro et al., 2016a; Thompson et al., 2018). Thus, the weights of the task-related and task-unrelated oscillations of these common synaptic inputs are determinants for precise force modulation (for a review, see Farina and Negro (2015)). Although the contributions of these oscillations to the generation of common synaptic input may not be directly measured in humans, they have been indirectly estimated through coherence analysis of discharge times of motor unit spike trains (Negro and Farina, 2012; Castronovo et al., 2015; Maillet et al., 2022; Rossato et al., 2022). For instance, Castronovo et al. (2015) have demonstrated significant coherence between motor unit spike trains up to approximately 80 Hz during isometric contractions, and these results have been repeatedly confirmed by subsequent studies using EMG (Kerkman et al., 2018) and motor unit (McManus et al., 2019; Muceli et al., 2022) recordings. Among these significant oscillation frequencies, the delta band of coherence (1-5 Hz) is believed to reflect the effective control signal to the motor neuron pool (i.e., task-related oscillations of the common synaptic input) as it encompasses the frequency range of the force signal (Farina et al., 2014). On the other hand, beta band oscillations (15-35 Hz) in motor unit coherence are commonly attributed to cortical origin, as studies have demonstrated associations between muscular and cortical activities within this frequency range (Conway et al., 1995b; Baker et al., 1997; Baker, 2007). Finally, the alpha band coherence (5-15 Hz) is typically associated with the physiological tremor (Conway et al., 1995a; Christakos et al., 2006; Laine et al., 2016). Given that alpha band frequencies are not entirely filtered out by the contractile properties of the muscle (Bawa and Stein, 1976; Baldissera et al., 1998), they are also reflected in the variability of force. Thus, these neural oscillations can be viewed as an involuntary common noise input that limits the accuracy of the force output (i.e., task-unrelated oscillations of the common synaptic input). For instance, a demanding visuomotor task has been shown to increase alpha band coherence, which was accompanied by larger force tremors (Laine et al., 2014). These findings, along with others (Ribot-Ciscar et al., 2009; Roche et al., 2011), suggest that a high-sensitivity task, such as the one presented to the participants in this study, would result in larger fluctuations in the alpha band and consequently, lead to increased force oscillations. Indeed, as illustrated in **Figure 2A** and reflected in the coefficient of variation of force results (**Figure 2B**), the *pre-skill acquisition* trial exhibited greater force fluctuations in both the TA and FDI muscles. Supporting this observation, we also found higher motor unit discharge variability, a measure of the variance of the common synaptic input received by the motor neurons, in the *pre-skill acquisition* trial (**Figures 3C** and **3D**), which is consistent with previous research (Knight and Kamen, 2004; Ely et al., 2022). However, we hypothesized that the repetition of this challenging task would prompt the central nervous system to effectively minimize the common synaptic oscillations unrelated to the specific task. This would lead to reductions in alpha band oscillations (physiological tremor), subsequent improvements in force control, and, ultimately, the short-term acquisition of the motor task.

To test our hypothesis, we experimentally examined changes in motor unit coherence within delta, alpha and beta bands between *pre-* and *post-skill acquisition*. For both investigated muscles, our results revealed reductions in z-coherence, specifically within the physiological tremor frequency band (**Figure 4**), supporting our hypothesis. Importantly, these reductions were reflected in the oscillations of muscle force output within the alpha band (**Figure 3C-E**). To further investigate whether these decreases in physiological tremor frequencies in both neural and force outputs were indeed associated with the force-matching skill acquisition, we calculated the linear coupling between the target and the force, as well as between the target and the CST (an estimation of the effective neural drive to the muscle). We found that both the target-force and target-CST coherence values increased significantly between *pre-* and *post-skill acquisition* (**Figure 5**), indicating an enhancement in the representation of task-related oscillations of shared synaptic input with the force-matching skill acquisition. Notably, this better match between the neural/mechanical oscillations and the target fluctuations correlated significantly with the observed reductions in alpha band (tremor) oscillations. These findings collectively support our hypothesis that, indeed, the short-term acquisition of a force-matching skill is mediated by a reduction of physiological tremor in motor neuron inputs.

The observed decrease in alpha band oscillations could be attributed to modulations in peripheral pathways, cortical pathways, or both. The possibility of peripheral modulation aligns with previous studies showing H-reflex depression following visuomotor skill tasks (Perez et al., 2005; Giboin et al., 2020), suggesting an increase in the presynaptic inhibition of Ia afferents during motor skill acquisition. Considering the involvement of Ia afferent loop in physiological tremor enhancement (Cresswell and Löscher, 2000; Christakos et al., 2006; Laine et al., 2016), this increased Ia inhibition with skill acquisition may underlie the observed reductions in physiological tremor oscillations (scenario B in **Figure 7**). Another possibility is that these decreases in tremor band occur via central pathways. This possibility is consistent with previous evidence showing that spinal interneurons could reduce alpha band oscillations in motor neuron output by phase-inverting the inputs to motor neurons at this frequency bandwidth (Williams et al., 2010; Koželj and Baker, 2014). These spinal interneurons would act as a neural filter, cancelling the descending oscillations within alpha band. Therefore, it is possible that the gain of this spinal interneuron filter undergoes upregulation during force-matching skill acquisition, consequently amplifying the cancellation effect (scenario A in **Figure 7**). To explore which of these two possibilities aligns more closely with our experimental outcomes, we conducted simulations. Our findings demonstrated that the spinal interneurons filter, canceling cortical oscillatory inputs in the alpha frequency range, is likely the explanation for our experimental findings, as all the results of this simulation scenario were similar to the experimental observations (**Figure 8**). Conversely, reductions in the gain of Ia afferent feedback loop alone are unlikely to explain the observed reductions in alpha band oscillations with the acquisition of the force-matching skill. However, it is still possible that more complex neural schemes of tremor oscillation cancellation occur, combining the two scenarios outlined in **Figure 7** into a unified system or, for example, involving recurrent inhibition from spinal Renshaw cells, which could also remove the alpha band components of motor neurons output (Williams and Baker, 2009).

A final consideration of the shared synaptic alterations during the learning of a force-matching task is related to the alterations in beta band oscillations. Our experimental results showed no significant changes in beta band coherence between *pre-* and *post-skill acquisition* trials for both FDI and TA muscles (**Figure 4**). Importantly, this lack of change was not attributed to a low number of tracked motor units between trials (see *Results*). However, in the simulation that best aligned with the experimental outcomes (scenario A in **Figure 7**), we could only replicate the experimental findings by simulating a slight increase in beta band oscillations in the shared synaptic inputs to motor neurons between *pre-* and *post-skill* acquisition models. These results suggest that beta oscillations might still play a role in short-term acquisition of a force-matching skill task. This finding aligns with recent evidence suggesting that cortical beta oscillations are associated, at least in part, with motor performance following visuomotor learning (Espenhahn et al., 2019). Further investigation is needed to elucidate the implications of beta oscillations in the context of a force-matching skill acquisition.

Finally, we observed a subtle increase in mean discharge rate (∼1 pps; **Figures 3A** and **3B**) with the force-matching skill acquisition. An increase in mean discharge rate has been linked to improved transmission of common synaptic input to motor neuron spike trains (Negro and Farina, 2012), which would result in increased shared synaptic oscillations. Contrarily, we observed a decrease in shared synaptic noise oscillations despite the increase in mean discharge rate. These results suggest that the observed increases in mean discharge rate are unlikely to explain the reductions in shared synaptic noise oscillations observed with force-matching skill acquisition. Hence, it is possible that changes in peripheral motor unit properties, such as reductions in motor unit twitch duration or amplitude, may have contributed to the observed changes in mean discharge rate. Another plausible possibility is the co-contraction of antagonistic muscles to stiffen the joints in response to the challenges of the learning task, inducing an increase in the mean discharge rate. Indeed, prior investigation has demonstrated that antagonistic co-contraction can enhance learning of novel motor tasks (Heald et al., 2018). However, further investigation is warranted to fully elucidate the underlying mechanisms.

In conclusion, our study demonstrates that the acquisition of a force-matching skill involves specific adaptations in the shared synaptic input to alpha motor neurons, leading to improved force control by minimizing physiological tremor oscillations in motor neuron inputs. Furthermore, our findings propose that the likely mechanism driving these reductions in alpha band oscillations involves spinal interneurons phase-cancelling the descending oscillations within alpha band. Therefore, our study provides novel insights into the neural mechanisms underpinning short-term motor learning.

## Materials and Methods

### Participants

Twenty-four healthy volunteers participated in the study. Specifically, two experiments were performed in which 13 participants (6 females; age: 31 ± 4 years; height: 176.8 ± 7.1 cm; mass: 71.9 ± 16.4 kg) had the FDI tested, and 11 (4 females; age: 31 ± 3 years; height: 174.9 ± 9.0 cm; mass: 71.2 ± 18.1 kg) the TA muscle. All participants were free from musculoskeletal or neurological injuries and provided written informed consent prior to the beginning of experiments. This study was approved by the local ethics committee (code NP5665) and conformed to the latest Declaration of Helsinki.

### Experimental design

Participants took part in a single experimental session lasting ∼1h. For the measurements on the FDI muscle, participants sat on an adjustable chair with their elbow flexed at 45° (0° being the anatomical position) and their right upper arm and hand comfortably resting on a custom-made device. The wrist and the hand were in a neutral position. The middle phalanx of the index finger was fixed to an adjustable support attached to a load cell (SM-100 N, Interface, Arizona, USA) so that the isometric abduction force produced by the index finger could be measured (**Figure 9A**). To standardize the hand position and minimize the contribution of other muscles, the little, ring, and middle fingers were separated from the index finger and secured to the device with Velcro straps. The forearm was also strapped to the device, and the thumb was secured at ∼80° angle to the index finger (**Figure 9A**). For the measurements on the TA muscle, participants were comfortably seated on a custom-built ankle dynamometer with their right knee fully extended, their ankle at 10° of plantar flexion (0° being the foot perpendicular to the shank), and their hip flexed at 70° (0° being the hip fully extended). The right foot was fixed with Velcro straps to an adjustable footplate perpendicularly connected to a load cell (SM-500 N, Interface, Arizona, USA) to record the dorsiflexion isometric force produced by the ankle (**Figure 9B**).

**Figure 9:**
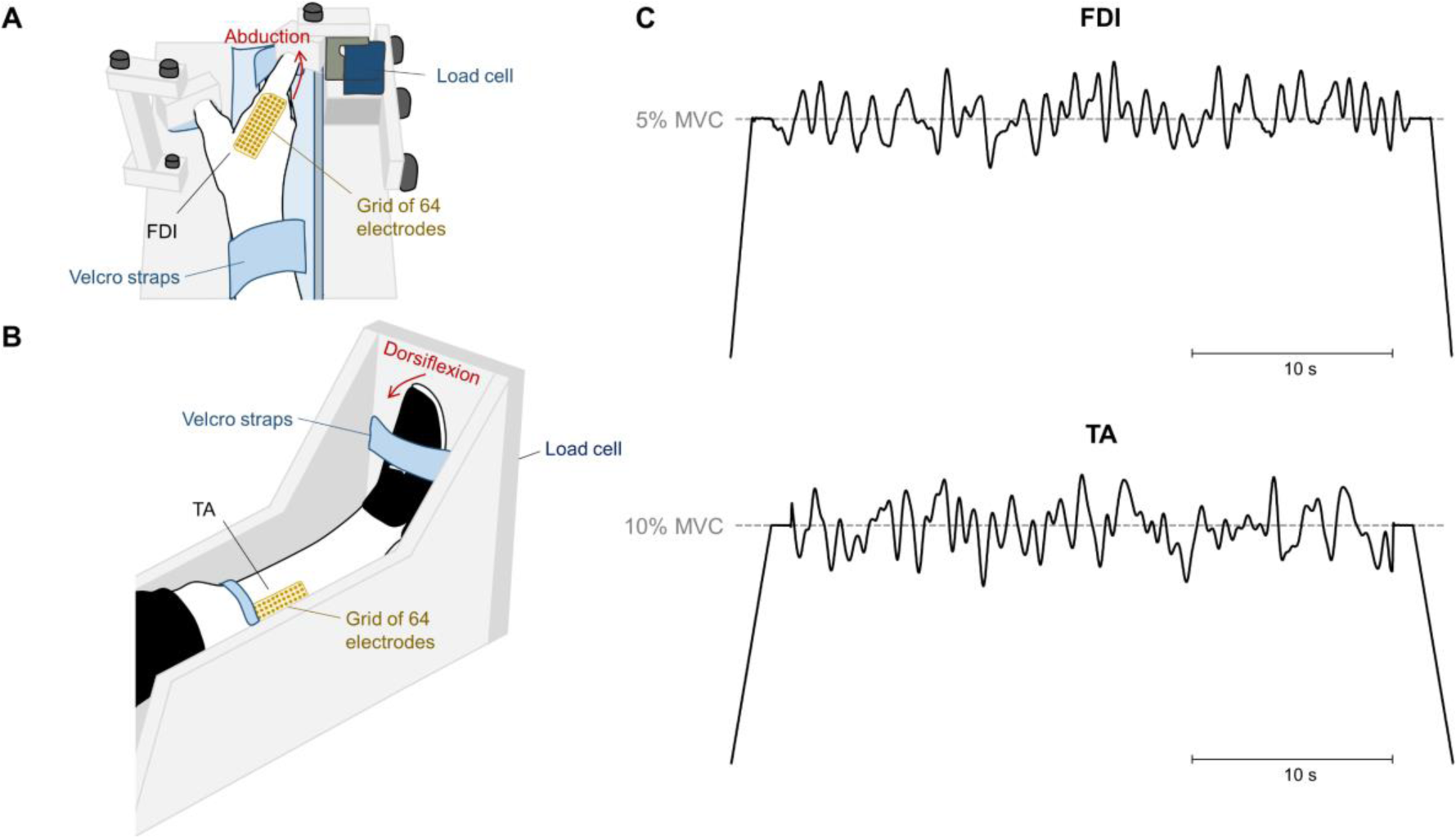
Experimental setup. A, position of the participants’ wrist and hand on the custom-made dynamometer to record the abduction isometric force produced by the index finger. B, position of the participants’ shank and foot on the custom-built dynamometer to measure the dorsiflexion isometric force produced by the ankle. High-density surface electromyography grids were placed over the first dorsal interosseous (FDI) muscle (A) and tibialis anterior (TA) muscle (B). C, Isometric force-matching tasks presented to the participants for the FDI (top) and the TA (bottom) muscles. Tasks involved a plateau region of 30 s containing a randomly generated signal low-pass filtered at 1.5 Hz. The target force level was defined as 5% and 10% of maximal voluntary contraction for the FDI and TA, respectively. The same trajectory was used throughout all trials of the force-matching skill task.

For both TA and FDI muscles, participants were initially asked to perform three isometric maximal voluntary contractions (MVCs) for 3 s, with a 60 s interval of rest in between. The greatest value across the three MVCs was considered the maximal isometric force and used as a reference for the following submaximal contractions. Then, after 5 min rest period, participants were instructed to perform 15 trials of a complex, isometric force-matching task (force-matching skill acquisition). The task involved a linear increase in force at a rate of 5% MVC/s, a variable force region for 30 s, and a linear decrease in force at a rate of 5% MVC/s. The variable force region contained oscillations above and below the target force level (averaged exerted force), which was defined as 5% MVC for the FDI and 10% MVC for the TA. The level of 5% MVC was chosen for the FDI to minimize fatigue effects across the task. Specifically, the oscillations consisted of a randomly generated signal with frequency content below 1.5 Hz (-3 dB low pass frequency). Two trajectories were generated, one for each muscle (**Figure 9C**), and the same trajectory was used throughout all trials and for all the subjects (Knight and Kamen, 2004). Each trajectory consisted of a black line, depicting the target force, over a white background and a red line depicting the subject’s force. A minimum of 60 s of rest was provided between trials, and prior to each trial, participants were encouraged by the same investigator to follow the target as closely as possible. Visual feedback from the target and the force was displayed on a computer monitor positioned at ∼60 cm in front of the participant, in which the entire target trajectory was visible and stationary.

### Data collection

During all trials of the force-matching task, HDsEMG signals were acquired from FDI and TA muscles using a grid of 64 electrodes arranged into 13 rows x 5 columns, with a missing electrode on a corner (4 mm inter-electrode distance for FDI; 8 mm inter-electrode distance for TA; OT Bioelettronica, Turin, Italy). An experienced investigator determined the position and orientation of the grids via palpation of anatomical landmarks. Specifically, the electrodes were attached over the belly of each muscle in the following locations: FDI, lateral to the line connecting the heads of the first and second metacarpals (**Figures 9A**); TA, ∼1 cm lateral to the tibial prominence (**Figure 9B**). Electrodes were fixed to the skin using a bi-adhesive foam, and the electrode-skin contact was ensured by filling the foam cavities with conductive paste (AC cream, Spes Medica, Genova, Italy). Reference electrodes were positioned on the right wrist for the FDI and on the right ankle for TA. Prior to electrode placement, the skin was cleaned with an abrasive paste (EVERI, Spes Medica, Genova, Italy) and shaved when necessary. Surface EMGs were recorded in monopolar mode and digitized at 2048 samples/s using a 16-bit amplifier (10-500 Hz bandwidth; Quattrocento, OT Bioelettronica, Turin, Italy). Force signals provided by the load cell were amplified by a factor of 100 (Forza-j, OT Bioelettronica, Turin, Italy) and sampled synchronously with EMGs.

### Data analysis

Force and HDsEMG signals were analyzed offline using MATLAB custom-written scripts.

*Force.* First, force signals were low-pass filtered at 15 Hz using a third-order Butterworth filter. Then, to quantify the performance for each trial of the force-matching task, the RMSE between the force and target signals was computed for the middle 30-s of the target (i.e., oscillatory region) (Knight and Kamen, 2004). Cross-correlation peak values were also calculated for the middle 30-s to assess similarities between fluctuations in force and target signals. For both RMSE and cross-correlation analyses, the detrended version of the signals was used. To assess force-matching improvements during the skill acquisition task, the first two and last two trials (out of the 15 trials) were selected for each participant. Since there were no statistical differences in RMSE and cross-correlation within both the initial two trials and the final two trials (see *Results* section), we chose to select one trial from the first two and one trial from the last two to represent the *pre-* and *post-skill acquisition* trials for all subsequent analysis. This decision was also driven by the aim to maximize the number of tracked motor units between trials (see *Identification and tracking of motor units*). Specifically, the trials selected were the ones with the highest and smallest RMSE between the force and target signals. In order to quantify how force steadiness changed during the learning task, we calculated the coefficient of variation (i.e., standard deviation/mean) of detrended force signals in the two selected trials. In addition, the power spectral density of force signals from the *pre-* and *post-skill acquisition* trials was estimated using Welch’s method (*pwelch* function in MATLAB; 1-s Hanning windows with 1948 samples of overlap). For each trial, the mean force power within the delta (1-5 Hz), and alpha (5-15 Hz) bands was calculated and retained for further analysis. Only the delta and alpha bands were considered in this analysis because the frequency range of force signals was up to 15 Hz.

*Identification and tracking of motor units.* Similar to the force, all motor unit analyses were performed for the middle 30-s of the target (i.e., oscillatory region). First, HDsEMG signals were bandpass filtered with a third-order Butterworth filter (20-500 Hz cut-off frequencies). After visual inspection, channels with low signal-to-noise ratio or artifacts were discarded. Then, the HDsEMG signals were decomposed into motor unit spike trains using a convolutive blind-source separation algorithm (Negro et al., 2016b). This method has been previously validated and extensively applied to assess the activity of single motor units (Castronovo et al., 2015; Negro et al., 2016b; Cogliati et al., 2020; Hassan et al., 2020). After the automatic identification of motor units, all the motor unit spike trains were visually inspected for false positives or false negatives (Hassan et al., 2020). Missing pulses or incorrectly assigned pulses producing non-physiological discharge rates were manually and iteratively edited by an experienced operator, and motor unit spike trains were re-estimated as previously proposed (Martinez-Valdes et al., 2017; Hassan et al., 2020). This approach has been shown to be highly reliable across operators (Hug et al., 2021). After the editing of motor unit spike trains, the motor units were tracked between the *pre-* and *post-skill acquisition* trials. This was achieved by reapplying the motor unit separation vectors, which are estimated with the blind source separation algorithm, from one trial to the other (Oliveira and Negro, 2021; Rossato et al., 2022). These motor unit separation vectors are unique for each individual motor unit and define the spatio-temporal matched filters to estimate the motor unit spike trains. This tracking procedure was performed in the forward and backward directions (i.e., motor unit separation vectors of *pre-skill acquisition* trial were applied on the *post-skill acquisition* trial, and vice-versa). Thus, our approach ensured that the same motor units were tracked and analyzed in both trials. Only motor units spike trains with a silhouette value, which is a metric to assess decomposition accuracy (Negro et al., 2016b), higher than 0.8 were used for analysis (Kapelner et al., 2020). For three participants in FDI and one participant in TA, we tracked the motor units based on their shape (Martinez-Valdes et al., 2017), as we were not able to track at least four motor units (see *Estimates of common synaptic input*) by reapplying the motor unit separation vectors. For this analysis, the two-dimensional representations of the motor unit action potentials of the identified motor units in the *pre-skill acquisition* trial were cross-correlated with the two-dimensional representations of the identified motor units in the *post-skill acquisition* trial (Martinez-Valdes et al., 2017; Cogliati et al., 2020). Only motor units with highly similar motor unit action potentials (cross-correlation > 0.8) were considered as belonging to the same motor units. The mean discharge rate and COV_ISI_ were calculated for each matched motor unit and stored for further analysis.

*Estimates of common synaptic input.* To assess changes in common synaptic input to motor neurons with the force-matching skill acquisition, coherence analysis was performed between CST of motor units from the same muscle (Negro and Farina, 2012; Castronovo et al., 2015; Negro et al., 2016a; Maillet et al., 2022; Rossato et al., 2022). Coherence is a frequency-domain linear coupling measure between two signals. This analysis was performed on two equally sized CSTs comprising motor units randomly selected from the identified, matched units. The number of motor units selected for each of the two groups was half of the total number of detected units, and this was repeated for up to 100 permutations. Only participants with at least four matched motor units were included in the coherence analysis. For all permutations, coherence was calculated between the two detrended CSTs using Welch’s periodogram with a 1-s Hanning window and an overlap of 95%. The obtained values were averaged for all permutations and transformed into standard z-scores as described by Gallet and Julien (2011). Only z-scores greater than the bias were considered for further analysis. The bias was determined as the mean value of z-scores between 250 and 500 Hz, as no significant coherence is expected in this frequency range (Maillet et al., 2022). To evaluate changes between *pre-* and *post-skill acquisition* trials, the areas under the curve of the z-coherence profiles within the delta (1-5 Hz), alpha (5-15 Hz) and beta (15-35 Hz) bands were calculated, and then, the area under the curve ratio (*post/pre*) was computed separately for each bandwidth. We subtracted one from the values of the area under the curve ratio so that values higher and lower than 0 indicated, respectively, an increase and decrease in z-coherence during *post-* compared with *pre-skill acquisition* trial. For purposes of visualization only, the pooled z-coherence across all participants was calculated as previously proposed (Baker et al., 2003).

Considering the nonlinear relation between the synaptic input to motor neuron and its output spike train becomes more accentuated at higher frequencies (due to the slow operating frequency of individual motor neurons; i.e., ∼30-40 pps), a greater number of motor units is required to accurately estimate coherence within the beta band (Farina and Negro, 2015). As detailed in the *Results* section, no significant differences were observed in z-coherence within this band between *pre-* and *post-skill acquisition* trials for both TA and FDI muscles. Therefore, to ensure that the absence of differences within this frequency band was not due to the number of motor units utilized in the CST, we conducted an additional analysis using the TA motor unit data (muscle with greater number of motor units matched between trials). Specifically, we calculated z-coherence between two CSTs, as described previously, but increasing the number of motor units in each CST incrementally from 1 to 13 (half of the maximal number of matched motor units across all participants). Subsequently, for each condition (1 motor unit in each CST, 2 motor units in each CST, and so forth), we computed the area under the curve ratio (*post/pre*) of z-coherence within the beta band. This process generated a curve depicting the z-coherence ratio within beta band as a function of the number of motor unit spike trains used in the CST. If our results within the beta band were influenced by the number of motor units used in the CSTs, we would expect to observe a significant deviation from zero in this curve as the number of motor units increased. Of note, in cases where a participant could not reach 13 motor units in each CST due to a lower number of identified units, we estimated the area under the curve of z-coherence within beta band by linear interpolation, using the other area under the curve values for that participant as reference points.

*Coherence between force/neural drive and target template.* To evaluate changes in the coupling between oscillations in force/neural drive and oscillations in the target template, we calculated the coherence between the force and the target template, and between the neural drive to muscles and the target template. In both cases, an increase in coherence values between *pre-* and *post-skill acquisition* trials would indicate an enhancement in the representation of shared synaptic input oscillations relevant to optimal force control in the intended motor task (i.e., task-related oscillations). For this analysis, the neural drive to the muscles was estimated by summing the binary discharge trains across all identified motor units (i.e., CST) (Thompson et al., 2018). Moreover, coherence was calculated only within the frequency bandwidth of the target template (i.e., delta band). Similar to motor unit analyses, the area under the curve ratio (*post/pre*) of z-coherence was computed to assess changes between *pre-* and *post-skill acquisition* trials.

### Simulations

To elucidate the neural mechanisms underlying the observed reductions in alpha band oscillations with the acquisition of the force-matching skill (see *Results*), we simulated the sequence of events from the excitation of an ensemble of motor neurons to the generation of isometric force output using the model proposed by Fuglevand et al. (1993). In this model, the population of motor neurons received common and independent inputs in varying relative proportions (Negro and Farina, 2012). Detailed descriptions of the modelling approach can be found in previous studies (Negro and Farina, 2011b; Farina et al., 2014; Negro et al., 2016a; Dideriksen and Negro, 2018). The motor neuron parameters were consistent with those used by Cisi and Kohn (2008) and were selected according to an exponential distribution over the pool of motor neurons (Fuglevand et al., 1993).

The number of motor neurons was set to 450 (similar to the number of TA motor units; Feinstein et al. (1955)), with only those having a minimum discharge rate of 8 pulses per second (pps) being fully recruited (Taylor et al., 2002). The input to the motor neuron pool was modelled as a linear summation of common synaptic input to all motor neurons and an independent noise input specific to each motor neuron (Negro et al., 2016a). The common synaptic input, which represents the input originating from the brain stem, spinal interneurons, or muscle afferents, included both task-related and task-unrelated oscillations. In our simulations, the task-related oscillations (i.e., oscillations within the delta band) were simulated using the same random signal provided as the target to participants during the skill acquisition task in the TA experimental recordings. Task-unrelated oscillations were simulated as the linear summation of 5-15 Hz Gaussian noise (i.e., oscillations within the alpha band) and 15-60 Hz Gaussian noise (i.e., oscillations within beta and piper bands). To explore the effects of afferent feedback modulation, we also included in the model a presynaptic gain of Ia afferent feedback into the motor neuron pool. Finally, the independent noise input, representing the individual variability of the membrane potential of each motor neuron, was modeled as a Gaussian noise with a bandwidth of 50 Hz (Negro et al., 2016a).

Two different scenarios were simulated to investigate the neural mechanisms underlying the observed changes between *pre-* and *post-skill acquisition* (refer to **Figure 7** in *Results*). In scenario A, we hypothesized that decreases in alpha band with the acquisition of the force-matching skill could be explained by spinal interneurons phase-cancelling central oscillatory inputs in the alpha frequency range (Williams et al., 2010; Koželj and Baker, 2014). The gain of this spinal interneurons filter would be upregulated during the skill acquisition task, thereby reducing alpha band oscillations between *pre-* and *post-skill acquisition*. To simulate this scenario, we created two models to represent *pre-* and *post-skill* acquisition. Then, we simulated an increase in the gain of the spinal interneurons filter in the *post-skill acquisition* model by decreasing the standard deviation of the 5-15 Hz input to the motor neuron pool compared with the *pre-skill acquisition* model. In scenario B, we hypothesized that reductions in alpha band with the force-matching skill acquisition could be explained by increases in presynaptic inhibition of Ia afferent feedback into the motor neuron pool. Similarly, we created two models to represent *pre-* and *post-skill* acquisition, but in this case, we simulated an increase in presynaptic inhibition of Ia afferent feedback in the *post-skill acquisition* model by decreasing the gain of the Ia afferent input to the motor neuron pool. In both scenarios, each model (*pre-* and *post-skill* acquisition) was repeated ten times as has been done in previous stimulation studies (Dideriksen and Negro, 2018). Following the same approach on the experimental data, we calculated the simulated force output power spectrum; the ratio (*post/pre*) of the area under the curve of motor unit z-coherence within delta (1-5 Hz), alpha (5-15 Hz) and beta (15-35 Hz) bands; and the area under the curve ratio (*post/pre*) of z-coherence between simulated force/CST and the target template. We then compared the results of scenarios A and B with the experimental results to explore which simulated scenario aligns more closely with the observed experimental outcomes.

### Statistical analysis

All statistical analyses were performed in R (version 4.3.0), using RStudio environment (version 2023.03.1).

To compare the RMSE between the four selected trials (first two and last two), Friedman tests were used. When significant effect of ‘trial’ was detected, post-hoc tests with Bonferroni correction were conducted for pairwise comparisons. To compare the coefficient of variation of force, and mean force power within delta and alpha bands between *pre-* and *post-skill acquisition* trials, Wilcoxon signed-rank tests were used. The Wilcoxon signed-rank test was also used to compare mean force power within delta and alpha bands between the *pre-skill acquisition* and *post-skill acquisition* models.

To compare mean discharge rate and COV_ISI_ between *pre-* and *post-skill acquisition* trials, linear mixed-effect models were applied, as they allow for the inclusion of all detected units and not just the mean value for each participant and trial (Boccia et al., 2019). This statistical model accounts for the non-independence of observations, which is particularly useful for this experimental design due to the hierarchical nature of motor unit data (greater correlation for units within participants compared with between participants; (Tenan et al., 2014)). For both mean discharge rate and COV_ISI_, random intercept models were applied with ‘trial’ (*pre-* and *post-skill acquisition*) as fixed effect and ‘participant’ as random effect (e.g., *mean discharge rate ∼ 1 + trial + (1 | participant)*). LMMs were implemented using the package *lmerTest* (Kuznetsova et al., 2017) with the Kenward-Roger method to approximate the degrees of freedom and estimate the *p*-values. The *emmeans* package was used to determine estimated marginal means and their differences with 95% confidence intervals (Lenth et al., 2019).

For both experimental and simulated data, to compare estimates of common synaptic input (i.e., z-coherence) between *pre-* and *post-skill acquisition* trials, changes in area under the curve ratio (*post/pre*) were tested using one-sample Wilcoxon signed-rank test (null hypothesis µ_0_ = 0), separately for delta, alpha and beta bands. To investigate whether the coherence results within the beta band were influenced by the number of motor units used in the CSTs, we utilized the one-sample Statistical Parametric Mapping test (Pataky et al., 2013). This test allowed us to determine whether the curve of the z-coherence ratio within beta band, as a function of the number of motor units (see details in *Estimates of common synaptic input*), showed a statistically significant deviation from 0. Conceptually, the one-sample Statistical Parametric Mapping test resembles the one-sample t-test, but it evaluates the entire curve. For both experimental and simulated data, changes in area under the curve ratio of target-force z-coherence and CST-force z-coherence between *pre-* and *post-skill acquisition* trials were assessed using the one-sample Wilcoxon signed-rank test (null hypothesis µ_0_ = 0).

For the experimental data, repeated measures correlations were performed to test whether changes in the motor unit z-coherence within alpha band were correlated with changes in z-coherence between force/neural drive and target template. Repeated measures correlations were implemented using the *rmcorr* package with fixed slopes to estimate a single correlation coefficient for all participants. For this analysis, the data of TA and FDI muscles were pooled together. For all statistical comparisons, statistical significance was set at an α of 0.05. For the results of motor unit discharge properties, the values in the text are reported as mean with 95% confidence intervals. All the other values are reported as mean ± standard deviation in the text and median/interquartile ranges in the figures. All individual data of motor unit discharge times for both TA and FDI muscles recorded in the *pre-* and *post-skill acquisition trials* are available at https://doi.org/10.6084/m9.figshare.23703804

## Acknowledgments

This study was funded by the European Research Council Consolidator Grant INcEPTION (contract no. 101045605).

## Author contributions

Conceptualization: H.V.C. and F.N.; Methodology: H.V.C., A.D.V. and F.N.; Software and Formal analysis: H.V.C. and F.N.; Investigation: H.V.C., A.C., A.B. and A.D.V.; Resources: C.O. and F.N.; Writing – original draft: H.V.C. and F.N.; Writing – review & editing: L.F., C.O., E.M.V. and F.N.; Supervision, project administration, and funding acquisition: F.N.

## References

Baker SN (2007) Oscillatory interactions between sensorimotor cortex and the periphery. Current Opinion in Neurobiology 17:649–655.

Baker SN, Olivier E, Lemon RN (1997) Coherent oscillations in monkey motor cortex and hand muscle EMG show task-dependent modulation. J Physiol 501 ( Pt 1):225–241.

Baker SN, Pinches EM, Lemon RN (2003) Synchronization in monkey motor cortex during a precision grip task. II. effect of oscillatory activity on corticospinal output. J Neurophysiol 89:1941–1953.

Baldissera F, Cavallari P, Cerri G (1998) Motoneuronal pre-compensation for the low-pass filter characteristics of muscle. A quantitative appraisal in cat muscle units. J Physiol 511 ( Pt 2):611–627.

Bawa P, Stein RB (1976) Frequency response of human soleus muscle. J Neurophysiol 39:788–793.

Boccia G, Martinez-Valdes E, Negro F, Rainoldi A, Falla D (2019) Motor unit discharge rate and the estimated synaptic input to the vasti muscles is higher in open compared with closed kinetic chain exercise. J Appl Physiol (1985) 127:950-958.

Castronovo AM, Negro F, Conforto S, Farina D (2015) The proportion of common synaptic input to motor neurons increases with an increase in net excitatory input. J Appl Physiol (1985) 119:1337-1346.

Christakos CN, Papadimitriou NA, Erimaki S (2006) Parallel neuronal mechanisms underlying physiological force tremor in steady muscle contractions of humans. J Neurophysiol 95:53–66.

Cisi RR, Kohn AF (2008) Simulation system of spinal cord motor nuclei and associated nerves and muscles, in a Web-based architecture. J Comput Neurosci 25:520–542.

Cogliati M, Cudicio A, Martinez-Valdes E, Tarperi C, Schena F, Orizio C, Negro F (2020) Half marathon induces changes in central control and peripheral properties of individual motor units in master athletes. J Electromyogr Kinesiol 55:102472.

Conway BA, Farmer SF, Halliday DM, Rosenberg JR (1995a) On the Relation between Motor-Unit Discharge and Physiological Tremor. In: Alpha and Gamma Motor Systems (Taylor A, Gladden MH, Durbaba R, eds), pp 596–598. Boston, MA: Springer US.

Conway BA, Halliday DM, Farmer SF, Shahani U, Maas P, Weir AI, Rosenberg JR (1995b) Synchronization between motor cortex and spinal motoneuronal pool during the performance of a maintained motor task in man. J Physiol 489 ( Pt 3):917–924.

Cresswell AG, Löscher WN (2000) Significance of peripheral afferent input to the alpha-motoneurone pool for enhancement of tremor during an isometric fatiguing contraction. Eur J Appl Physiol 82:129–136.

Dayan E, Cohen LG (2011) Neuroplasticity subserving motor skill learning. Neuron 72:443–454.

Dideriksen JL, Negro F (2018) Spike-triggered averaging provides inaccurate estimates of motor unit twitch properties under optimal conditions. J Electromyogr Kinesiol 43:104–110.

Ely IA, Jones EJ, Inns TB, Dooley S, Miller SBJ, Stashuk DW, Atherton PJ, Phillips BE, Piasecki M (2022) Training-induced improvements in knee extensor force accuracy are associated with reduced vastus lateralis motor unit firing variability. Experimental Physiology 107:1061–1070.

Enoka RM, Farina D (2021) Force Steadiness: From Motor Units to Voluntary Actions. Physiology 36:114–130.

Espenhahn S, van Wijk BCM, Rossiter HE, de Berker AO, Redman ND, Rondina J, Diedrichsen J, Ward NS (2019) Cortical beta oscillations are associated with motor performance following visuomotor learning. Neuroimage 195:340–353.

Evans CM, Baker SN (2003) Task-dependent intermanual coupling of 8-Hz discontinuities during slow finger movements. Eur J Neurosci 18:453–456.

Farina D, Negro F (2015) Common synaptic input to motor neurons, motor unit synchronization, and force control. Exerc Sport Sci Rev 43:23–33.

Farina D, Negro F, Dideriksen JL (2014) The effective neural drive to muscles is the common synaptic input to motor neurons. J Physiol 592:3427–3441.

Feinstein B, Lindegard B, Nyman E, Wohlfart G (1955) Morphologic studies of motor units in normal human muscles. Acta Anat (Basel) 23:127–142.

Fuglevand AJ, Winter DA, Patla AE (1993) Models of recruitment and rate coding organization in motor-unit pools. J Neurophysiol 70:2470–2488.

Gallet C, Julien C (2011) The significance threshold for coherence when using the Welch’s periodogram method: Effect of overlapping segments. Biomedical Signal Processing and Control 6:405–409.

Giboin L-S, Tokuno C, Kramer A, Henry M, Gruber M (2020) Motor learning induces time-dependent plasticity that is observable at the spinal cord level. The Journal of Physiology 598:1943–1963.

Halliday AM, Redfearn JW (1956) An analysis of the frequencies of finger tremor in healthy subjects. J Physiol 134:600–611.

Hassan A, Thompson CK, Negro F, Cummings M, Powers RK, Heckman CJ, Dewald JPA, McPherson LM (2020) Impact of parameter selection on estimates of motoneuron excitability using paired motor unit analysis. J Neural Eng 17:016063.

Heald JB, Franklin DW, Wolpert DM (2018) Increasing muscle co-contraction speeds up internal model acquisition during dynamic motor learning. Scientific Reports 8:16355.

Hug F, Avrillon S, Del Vecchio A, Casolo A, Ibanez J, Nuccio S, Rossato J, Holobar A, Farina D (2021) Analysis of motor unit spike trains estimated from high-density surface electromyography is highly reliable across operators. J Electromyogr Kinesiol 58:102548.

Jensen JL, Marstrand PC, Nielsen JB (2005) Motor skill training and strength training are associated with different plastic changes in the central nervous system. J Appl Physiol (1985) 99:1558-1568.

Johansson RS, Westling G (1988) Programmed and triggered actions to rapid load changes during precision grip. Exp Brain Res 71:72–86.

Kapelner T, Sartori M, Negro F, Farina D (2020) Neuro-Musculoskeletal Mapping for Man-Machine Interfacing. Scientific Reports 10:5834.

Karni A, Meyer G, Jezzard P, Adams MM, Turner R, Ungerleider LG (1995) Functional MRI evidence for adult motor cortex plasticity during motor skill learning. Nature 377:155–158.

Karni A, Meyer G, Rey-Hipolito C, Jezzard P, Adams MM, Turner R, Ungerleider LG (1998) The acquisition of skilled motor performance: Fast and slow experience-driven changes in primary motor cortex. Proceedings of the National Academy of Sciences 95:861–868.

Kerkman JN, Daffertshofer A, Gollo LL, Breakspear M, Boonstra TW (2018) Network structure of the human musculoskeletal system shapes neural interactions on multiple time scales. Sci Adv 4:eaat0497.

Kleim JA, Barbay S, Nudo RJ (1998) Functional reorganization of the rat motor cortex following motor skill learning. J Neurophysiol 80:3321–3325.

Knight CA, Kamen G (2004) Enhanced motor unit rate coding with improvements in a force-matching task. Journal of Electromyography and Kinesiology 14:619–629.

Koželj S, Baker SN (2014) Different phase delays of peripheral input to primate motor cortex and spinal cord promote cancellation at physiological tremor frequencies. J Neurophysiol 111:2001–2016.

Kumar A, Tanaka Y, Grigoriadis A, Grigoriadis J, Trulsson M, Svensson P (2017) Training-induced dynamics of accuracy and precision in human motor control. Scientific Reports 7:6784.

Kuznetsova A, Brockhoff PB, Christensen RHB (2017) lmerTest Package: Tests in Linear Mixed Effects Models. Journal of Statistical Software 82:1–26.

Laine CM, Yavuz SU, Farina D (2014) Task-related changes in sensorimotor integration influence the common synaptic input to motor neurones. Acta Physiol (Oxf) 211:229–239.

Laine CM, Nagamori A, Valero-Cuevas FJ (2016) The Dynamics of Voluntary Force Production in Afferented Muscle Influence Involuntary Tremor. Front Comput Neurosci 10:86.

Lenth R, Singmann H, Love J, Buerkner P, Herve M (2019) Package ‘emmeans’. In. Lippold O (1971) Physiological tremor. Sci Am 224:65-73.

Maillet J, Avrillon S, Nordez A, Rossi J, Hug F (2022) Handedness is associated with less common input to spinal motor neurons innervating different hand muscles. J Neurophysiol 128:778–789.

Marsden CD, Meadows JC, Lange GW, Watson RS (1967) Effect of deafferentation on human physiological tremor. Lancet 2:700–702.

Martinez-Valdes E, Negro F, Laine CM, Falla D, Mayer F, Farina D (2017) Tracking motor units longitudinally across experimental sessions with high-density surface electromyography. J Physiol 595:1479–1496.

McAuley JH, Marsden CD (2000) Physiological and pathological tremors and rhythmic central motor control. Brain 123 ( Pt 8):1545–1567.

McAuley JH, Rothwell JC, Marsden CD (1997) Frequency peaks of tremor, muscle vibration and electromyographic activity at 10 Hz, 20 Hz and 40 Hz during human finger muscle contraction may reflect rhythmicities of central neural firing. Exp Brain Res 114:525-541.

McManus L, Flood MW, Lowery MM (2019) Beta-band motor unit coherence and nonlinear surface EMG features of the first dorsal interosseous muscle vary with force. J Neurophysiol 122:1147–1162.

Muceli S, Poppendieck W, Holobar A, Gandevia S, Liebetanz D, Farina D (2022) Blind identification of the spinal cord output in humans with high-density electrode arrays implanted in muscles. Science Advances 8:eabo5040.

Negro F, Farina D (2011a) Linear transmission of cortical oscillations to the neural drive to muscles is mediated by common projections to populations of motoneurons in humans. J Physiol 589:629–637.

Negro F, Farina D (2011b) Decorrelation of cortical inputs and motoneuron output. J Neurophysiol 106:2688–2697.

Negro F, Farina D (2012) Factors influencing the estimates of correlation between motor unit activities in humans. PLoS One 7:e44894.

Negro F, Holobar A, Farina D (2009) Fluctuations in isometric muscle force can be described by one linear projection of low-frequency components of motor unit discharge rates. J Physiol 587:5925–5938.

Negro F, Yavuz U, Farina D (2016a) The human motor neuron pools receive a dominant slow-varying common synaptic input. J Physiol 594:5491–5505.

Negro F, Muceli S, Castronovo AM, Holobar A, Farina D (2016b) Multi-channel intramuscular and surface EMG decomposition by convolutive blind source separation. J Neural Eng 13:026027.

Nudo RJ, Milliken GW, Jenkins WM, Merzenich MM (1996) Use-dependent alterations of movement representations in primary motor cortex of adult squirrel monkeys. J Neurosci 16:785–807.

Oliveira AS, Negro F (2021) Neural control of matched motor units during muscle shortening and lengthening at increasing velocities. J Appl Physiol (1985) 130:1798-1813.

Pascual-Leone A, Tarazona F, Catalá MD (1999) Applications of transcranial magnetic stimulation in studies on motor learning. Electroencephalogr Clin Neurophysiol Suppl 51:157–161.

Pascual-Leone A, Nguyet D, Cohen LG, Brasil-Neto JP, Cammarota A, Hallett M (1995) Modulation of muscle responses evoked by transcranial magnetic stimulation during the acquisition of new fine motor skills. J Neurophysiol 74:1037–1045.

Pataky TC, Robinson MA, Vanrenterghem J (2013) Vector field statistical analysis of kinematic and force trajectories. Journal of Biomechanics 46:2394–2401.

Perez MA, Lungholt BKS, Nielsen JB (2005) Presynaptic control of group Ia afferents in relation to acquisition of a visuo-motor skill in healthy humans. The Journal of Physiology 568:343–354.

Perez MA, Lungholt BK, Nyborg K, Nielsen JB (2004) Motor skill training induces changes in the excitability of the leg cortical area in healthy humans. Exp Brain Res 159:197–205.

Plautz EJ, Milliken GW, Nudo RJ (1995) Differential effects of skill acquisition and motor use on the reorganization of motor representations in area 4 of adult squirrel monkeys.. In: Society for Neuroscience.

Ribot-Ciscar E, Hospod V, Roll JP, Aimonetti JM (2009) Fusimotor drive may adjust muscle spindle feedback to task requirements in humans. J Neurophysiol 101:633–640.

Roche N, Bussel B, Maier MA, Katz R, Lindberg P (2011) Impact of precision grip tasks on cervical spinal network excitability in humans. The Journal of Physiology 589:3545–3558.

Rossato J, Tucker K, Avrillon S, Lacourpaille L, Holobar A, Hug F (2022) Less common synaptic input between muscles from the same group allows for more flexible coordination strategies during a fatiguing task. J Neurophysiol 127:421–433.

Taylor AM, Steege JW, Enoka RM (2002) Motor-unit synchronization alters spike-triggered average force in simulated contractions. J Neurophysiol 88:265–276.

Tenan MS, Marti CN, Griffin L (2014) Motor unit discharge rate is correlated within individuals: a case for multilevel model statistical analysis. J Electromyogr Kinesiol 24:917–922.

Thompson CK, Negro F, Johnson MD, Holmes MR, McPherson LM, Powers RK, Farina D, Heckman CJ (2018) Robust and accurate decoding of motoneuron behaviour and prediction of the resulting force output. J Physiol 596:2643–2659.

Ungerleider LG, Doyon J, Karni A (2002) Imaging brain plasticity during motor skill learning. Neurobiol Learn Mem 78:553–564.

Williams ER, Baker SN (2009) Renshaw cell recurrent inhibition improves physiological tremor by reducing corticomuscular coupling at 10 Hz. J Neurosci 29:6616–6624.

Williams ER, Soteropoulos DS, Baker SN (2010) Spinal interneuron circuits reduce approximately 10-Hz movement discontinuities by phase cancellation. Proceedings of the National Academy of Sciences 107:11098–11103.

Willingham DB (1998) A neuropsychological theory of motor skill learning. Psychol Rev 105:558–584.

